# Comprehensive Assessment of Eleven *de novo* HiFi Assemblers on Complex Eukaryotic Genomes and Metagenomes

**DOI:** 10.1101/2023.06.29.546998

**Authors:** Wenjuan Yu, Haohui Luo, Jinbao Yang, Shengchen Zhang, Heling Jiang, Xianjia Zhao, Xingqi Hui, Da Sun, Liang Li, Xiu-qing Wei, Stefano Lonardi, Weihua Pan

## Abstract

**Background:** Pacific Bioscience HiFi sequencing technology generates long reads (>10 kbp) with very high accuracy (less than 0.01% sequencing error). While several *de novo* assembly tools are available for HiFi reads, there are no comprehensive studies on the evaluation of these assemblers.

**Results:** We evaluated the performance of eleven *de novo* HiFi assemblers on (i) real data for three eukaryotic genomes, (ii) 34 synthetic datasets with different ploidy, sequencing coverage levels, heterozygosity rates and sequencing error rates, (iii) one real metagenomic dataset, and (iv) five synthetic metagenomic datasets with different composition abundance and heterozygosity rates. The nine assemblers were evaluated using QUAST (Quality Assessment Tool) and BUSCO (Benchmarking Universal Single-Copy Ortholog). We also used several additional criteria, namely, completion rate, single-copy completion rate, duplicated completion rate, average proportion of largest category, average distance difference, quality value, run-time and memory utilization. On complex eukaryotic genomes, Hifiasm had a clear advantage over the other assemblers in all tested experiments. On synthetic datasets, Hifiasm, HiCanu, and HiFlye performed equally well. Shasta and Peregrine had good performance across varying ploidy, but required high computational resources. On metagenomic datasets, Hifiasm-meta demonstrated a clear advantage over other assemblers.

**Conclusion:** We carried out a comprehensive benchmarking study of commonly used assemblers on complex eukaryotic genomes and metagenomes. Our study will help the research community to choose the most appropriate assembler for their data and identify possible improvements in assembly algorithms.

## INTRODUCTION

Advances in sequencing technology have been a driving force in molecular biology and genomics, in particular for *de novo* genome assembly (1-3). Single molecule sequencing (SMS) technologies recently introduced on the market can generate longer reads that can span longer genomic repeats, thus have reduced the computational cost and simplified the *de novo* assembly problem. A notable example of SMS is Pacific Biosciences HiFi technology that can provide long reads (>10 kbp) with very high accuracy (less than 0.01% sequencing error). In contrast Illumina sequencing instruments generate very high volumes of much shorter (100-250 bp) and accurate reads (less than 1% sequencing error); Oxford Nanopore generates very long reads (up to 1.5 Mbp) but sequencing error rate can be as high as 15%. PacBio HiFi reads have enabled significant improvements in the assembly of the human genome (4,5) as well several other eukaryotic genomes (6-11).

The problem of *de novo* assembly can be computationally challenging because of the high repetitive content of genomes, sequencing errors, non-uniform or insufficient sequencing coverage and chimeric reads. In the literature, the problem is solved either using the overlap-layout consensus (OLC) (12), or the de Bruijin graph (DBG) (12,13), or the string graph (14), depending on the nature and the number of the input reads. These methods also play an essential role in SMS assembly (6).

Assemblers for long SMS reads also use OLC, DBG and the string graph (or any combination thereof). For instance, Canu (15) integrates hybrid error correction PBcR (16), MinHash Alignment Process (MHAP), and some modules from the Celera Assembler (17). miniasm (18) implements the overlap and layout steps in the OLC assembly paradigm, but not the consensus. The absence of the last step makes miniasm particularly fast, but its application is limited to relatively small genomes that are not very repetitive (18). HiCanu (19) is a special version of Canu that leverages the high-quality of HiFi reads. FALCON (20) builds primary contigs via a string graph, and then generates the haplotype-resolved assembly using phased reads. Shasta was designed for the assembly of human genome with Oxford Nanopore reads, but the developers have recently added a HiFi-mode (21). Peregrine (22) takes advantage of the high accuracy of HiFi reads and uses sparse hierarchical minimizers to index reads thereby reducing the high computational cost of all-against-all alignment step in the OLC pipeline. hifiasm was specifically designed for HiFi reads: it uses a phased assembly graph to reconstruct the haplotype of diploid genomes (23). hifiasm-meta represents a variant of hifiasm devised to carry out the assembly of multiple genomes present within a metagenomic sample. This tool incorporates a novel read selection step and introduces innovative criteria aimed at safeguarding reads that originate from genomes with limited coverage (24). ABruijn (25), Flye (26), HiFlye, and metaFlye (27) are based on the de Bruijn graph. ABruijn combines DBG and the OLC approaches. Flye uses the repeat graph, which extends the de Bruijn Graph so that it can deal with the errors in long reads. wtdbg2 also uses a combination of OLC and DBG (28): it utilizes a data structure called fuzzy-Bruijn graph to enable an efficient all-versus-all read alignment. MECAT and NECAT are assemblers for PacBio reads and ONT reads, respectively, that use a fast-scoring method to filter out spurious alignments (29,30). Verkko is an improved Canu assembler that can assemble HiFi reads and ONT reads simultaneously and was used for the telomere-to-telomere assembly of the human genome (31).

At the time of writing, users can choose from at least twenty-five assemblers for SMS reads (see **Supplementary Table 1**) depending on the type of reads they have (15,18-36). However, choosing the “best assembler” for their data is a daunting proposition, because the performance of an assembler depends on organism ploidy, genome repetitive content, genome size, heterozygosity, and many other factors. Even if we focus only on HiFi assemblers, there is no comprehensive study that could guide users on the expected performance of these assemblers on large eukaryotic genomes. The only studies we could find were (1) Zhang et al. (37), who tested Flye, HiCanu, hifiasm, NECAT and NextDenovo on HiFi and ONT reads, but only on brewer’s yeast; (2) Gavrielatos et al. (38), who tested Canu, hifiasm, WENGAN and HiCanu on Hifi and ONT reads, but only on the fruit fly and human haploid genome, and in the context of hybrid assembly.

To address this shortcoming, here we report on a comprehensive assessment of the performance of eleven SMS assemblers for HiFi reads (nine for genome assembly and two for metagenomes). Our choice of these eleven assemblers was based on their (i) popularity (based on their usage/citations), (ii) user friendliness (e.g., how easy it is to install and run them), (iii) algorithmic novelty, and (iv) the fact that they are currently actively maintained. The eleven HiFi assemblers we selected are HiCanu, hifiasm, HiFlye, hifiasm-meta, metaFlye, Peregrine, Shasta, Verkko, MECAT2, miniasm, and NextDenovo. We studied the performance of these assemblers under various conditions, including various sequencing coverage, heterozygosity, and ploidy (see **Supplementary Table 2**). The eleven assemblers were evaluated using QUAST/ MetaQUAST (39,40) (Quality Assessment Tool) and BUSCO (Benchmarking Universal Single-Copy Ortholog). We also measured new quality metrics, namely completion rate, single-copy completion rate, duplicated completion rate, average proportion of largest category, average distance difference, and quality value. We also recorded run-time and memory utilization.

## MATERIALS AND METHODS

### Datasets

#### Real datasets with varying ploidy

We tested the performances of the assemblers on the real HiFi reads from three eukaryotic genomes: (i) the homozygous diploid rice (*Oryza sativa*) genome ZS97 (9), (ii) the heterozygous diploid potato (*Solanum tuberosum*) genome (41) (NCBI project PRJNA686812 and PRJNA573826) and (iii) the autotetraploid wax apple (*Syzygium samarangense*) genome. The HiFi reads for rice and potato were downloaded from NCBI, whereas those for the wax apple were generated as part of this study (NCBI project PRJNA928838) due to the lack of HiFi data for polyploid genomes in NCBI. The ‘Tub’ variety of wax apple was selected for sequencing: young leaves were collected from an individual tree planted in the field of Fujian Academy of Agricultural Sciences (Fujian province, China) under the voucher number GPLWFJGSS0058. Genomic DNA was isolated using the QIAGEN Plant Genomic DNA Kit according to the standard PacBio operating procedure. Then, the genomic DNA was sheared by g-TUBE (Covaris, Woburn, MA, USA) resulting in 6 kb-20 kb fragments, and sequenced using a PacBio Sequel II instrument using the HiFi protocol, generating 39x coverage. Two of the authors of this paper have previously generated PacBio CLR, ONT and Hi-C data for the wax apple for another sequencing project (42). With the addition of the HiFi reads, the wax apple dataset is now the most comprehensive dataset for polyploid genomes. The statistics of these datasets are shown in **Supplementary Table 3**.

#### Synthetic datasets with varying ploidy

To carry out a comprehensive evaluation of the performance of the HiFi assemblers, the homozygous diploid rice genome ZS97 was used to produce (i) a synthetic heterozygous diploid genome and (ii) a synthetic autotetraploid genome. First, we generated two sets of synthetic chromosomes (four in the case of the autotetraploid) by adding SNPs and structural variations (insertions and deletions) to the ZS97 genomes (9) using a custom script (https://github.com/sc-zhang/bioscripts/blob/master/SimSID.py). To generate realistic synthetic genomes, we tuned the parameters of SimSID.py until the reads generated by wigsim (https://github.com/lh3/wgsim) from the synthetic genomes produced a GenomeScope2 *k*-mer distribution that matched the *k*-mer distribution of the heterozygous diploid big berry manzanita (43), and the autotetraploid potato (44), respectively (**Supplementary Figure 1**) (43,44). Following this procedure the synthetic heterozygous diploid genome was generated using a heterozygosity rate of 2.6%, while the autotetraploid genome was generated using a heterozygosity rate of 1.0%. The proportion of SNPs, insertions and deletions were 1:1:1. PBsim v1.0.4 (45) was used to generate synthetic HiFi reads for (i) the ZS97 genome (homozygous diploid), (ii) the synthetic heterozygous diploid genome, and (iii) the synthetic autotetraploid genome. Detailed statistics for these datasets are shown in **Supplementary Table 4**.

#### Synthetic datasets with varying heterozygosity rates, coverage levels and sequencing error rates

To generate synthetic datasets with varying heterozygosity rates, we generate four diploid genomes with heterozygosity rates 0.5%, 1.0%, 1.5% and 2.5%. The proportion of SNPs, insertions and deletions was 2:1:1(parameter is recommended in (46)). HiFi reads were simulated by PBsim v1.0.4. Detailed statistics for these datasets are shown in **Supplementary Table 5**.

To generate the synthetic datasets with varying read coverages, we used PBsim v1.0.4 on the synthetic heterozygous diploid genome with different coverages by changing the parameter “--depth”. Detailed statistics for these datasets are shown in **Supplementary Table 6**.

To generate the synthetic datasets with varying sequencing error rates, we used PBsim v1.0.4 to simulate HiFi reads for the ZS97 genome and the two synthetic genomes in with coverage 20x and the sequencing error models trained on the real HiFi datasets shown in **Supplementary Table 7**. Detailed statistics of these synthetic datasets are shown in **Supplementary Table 8**.

#### Real and synthetic metagenome datasets

To evaluate the performance of metagenome assemblers, we utilized a real sheep gut dataset (NCBI project PRJNA595610) (47) and created five synthetic datasets. Detailed statistics of the sheep gut dataset are shown in **Supplementary Table 9**.

To build the first synthetic datasets, we selected 382 bacterial genomes from the GTDB database covering 90 genera of Bacteroidetes, Actinobacteria, Firmicutes, Proteobacteria, and Fusobacteria phyla, and merged them into a metagenome by assigning different abundances to them. The abundance profile was obtained by sampling coverage values from a real chicken gut metagenome assembly (48). PBsim v1.0.4 was used to simulate the HiFi reads with the error model learned on the real sheep gut dataset. Detailed statistics for this synthetic dataset are shown in **Supplementary Table 10**.

The other four synthetic datasets were built for different taxonomic levels such as Phylum, Class, Family and Genus. For example, to build the Phylum dataset, 200 genomes from the same Phylum were selected, and abundances and the simulated HiFi reads were generated in the same way as the first synthetic datasets. The synthetic datasets of Class, Family and Genus were also generated in the same way as Phylum. Detailed statistics for these synthetic datasets are shown in **Supplementary Table 11**.

### Assembler performance evaluation

Our comprehensive assessment of the assemblers was carried out by evaluating completeness, contiguity, and accuracy of assembled genomes for real and synthetic data sets.

#### Contiguity

QUAST v5.0.2 was used to assess the contiguity of the assemblies. We recorded N50, L50, and the length of the longest contig. We recall that the N50 is defined as the length for which the set of all contigs of that length or longer covers at least half of the assembled genome. L50 is defined as the count of the smallest number of contigs whose total length makes up half of the assembled genome. NG50 is defined as the length for which the set of all contigs of that length or longer covers at least 50% of the length of the actual genome. LG50 is defined as the count of the smallest number of contigs whose total length makes up half of the actual genome.

#### Completeness

BUSCO (Benchmarking Universal Single-Copy Orthologs) was used to assess genome completeness on all assemblies produced on real data. BUSCO measures the fraction of highly conserved genes that are present in the assembly (full length or fragmented).

On simulated data, three additional criteria were used to measure completeness, namely *completeness rate* (CR), *single-copy completeness rate* (SCR), and *duplicated completeness rate* (DCR). CR, SCR and DCR are based on *k*-mer analysis and are explained next with the help of **Figure 1**. In all our experiments we used *k* = 21, which is the value typically used for eukaryotic genomes (49). We call the set of *k*-mers that are unique in the reference genome *S*_*RefG*_*unikmers*_, and the set of all *k*-mers in the genome assembly *S*_*CtgG*_*kmers*_ (refer to **Figure 1**). If the assembly is complete, then all unique *k*-mers in the reference are expected to appear in the assembly. In general, however *S*_*RefG*_*unikmers*_ is a subset of *S*_*CtgG*_*kmers*_. Thus, we define the *completeness rate* (CR) as follows

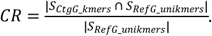

CR ranges between 0 and 1, where 1 indicates a complete assembly.

Observe that the set 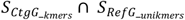 contains single copy *k*-mers and duplicated *k*-mers. Thus, we can define the *single-copy completeness rate* (SCR) as follows

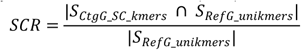

where 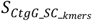 is the set of single copy *k*-mers in the assembly. Similarly, the *duplicated completeness* rate (DCR) is defined as

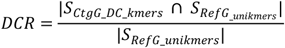

where *S*_*CtgG_DC_kmers*_ is the set of duplicated *k*-mers in the assembly. Obviously, *CR* = *SCR* + *DCR*.

**Figure 1.**
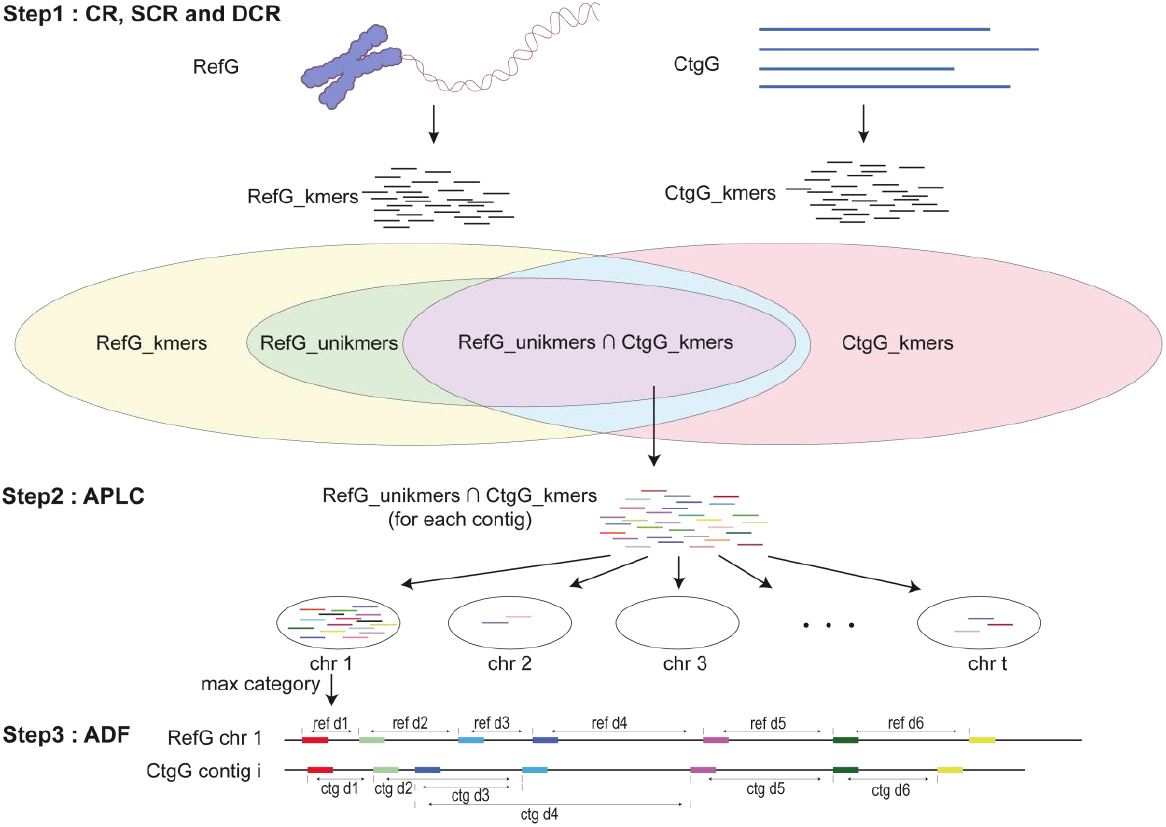
Completeness and accuracy of assembled genomes on synthetic data are evaluated based on a *k*-mer analysis. **Step1:** The completeness rate (CR), single-copy completeness rate (SCR), and duplicated completeness rate (DCR) are computed from the shared *k*-mers that are unique in the reference and the *k*-mers in the assembled contigs (represented in purple). **Step2**: The average proportion of the largest category (APLC) is computed from the *k*-mers in common between the unique k-mers in the reference genome and *k*-mers in specific contigs. **Step3**: The average distance difference (ADF) is computed from pairs of adjacent unique *k*-mers.

#### Accuracy

To evaluate the assembly accuracy, we use the *consensus quality value* (QV), the *N-rate*, the *average proportion of the largest category* (APLC), and the *average distance difference* (ADF).

The *consensus quality value* (QV) was defined in (49), and it measures the frequency of consensus errors in the assembly. QV is defined as follows

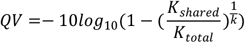

where the *K*_*total*_ is the total number of *k*-mers found in an assembly and_*shared*_ is the number of shared *k*-mers between the assembly and the reads. Observe that 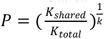 represents the probability that a base in the assembly is correct, and −10*l*og_10_(1 − P) can be interpreted as a Phred quality score (50). For instance, QV>60 indicates an excellent assembly at the base-level (for instance, QV=60 translates to an accuracy of 99.9999%).

The *N-rate* is the proportion of ambiguous bases (‘N’s) in the assembly, as reported by QUAST. The lower is the N-rate, the better the assembly.

Now observe that a contig mis-assembly can be detected if that contig contains unique *k*-mers from two or more chromosomes. We proposed a measure called the *average proportion of the largest category* (APLC), which captures the mis-assembly rate for each assembled contigs. First, we define *Pchr*_*a,c*_, which is the proportion of unique *k*-mers for chromosome *a* that appear in contig *c*, as follows

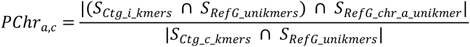

where *a in* [1, t], *t* is the number of chromosomes, *S*_*RefG*_*unikmers*_is the set of unique *k*-mers in chromosome *a*, and *S*_*Ctg*_*c*_*kmers*_ is the set of *k*-mers in contig *c*. Next, we define *PChr*_*c*_ as the largest value of *PChr*_*c,a*_ over all the t chromosomes for contig *c*, as follows

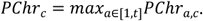

Finally, we can define the *average proportion of the largest category* (APLC) is the average value *PL*_*c*_ over all the contigs, defined as follow

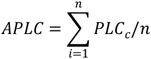

where *n* is the number of the contigs in the assembly.

The final measure of accuracy is based on the distance between pairs of unique adjacent *k*-mer, which is expected to be the same in the reference genome and the assembled contigs. First, unique *k*-mers are sorted by their position in the reference genome, then the distance of those *k*-mers in the assembly is calculated. The difference *DF*_*c*_ for a contig *c*, is defined as follows

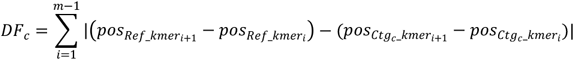

where *m* is the number of unique *k*-mer, 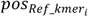 is the position of the *i*-th *k*mer in the reference genome, and 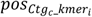 is the position of the *i*-th *k*-mer in assembled contig *c*. Finally, we can define the *average distance difference* (ADF) as follows

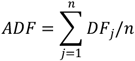

where *n* is the number of the contigs in the assembly. The smaller the value of ADF, the more accurate the assembly is.

### Runtime and memory usage

Time and memory usage were recorded for all experiments in this study. All assemblers were run on an Inspur Cluster Engine Linux cluster at Agricultural Genomics Institute at Shenzhen, Chinese Academy of Agricultural Sciences. The cluster has six main nodes, each of which has 80 CPUs and 3TB memory.

## RESULTS

A comprehensive set of experiments were conducted on the eleven genome assemblers using both real and simulated data for various choices of ploidy. Four assemblers were selected for a deeper analysis on datasets produced for different choices of sequencing coverage and heterozygosity. Finally, four metagenome assemblers were tested using both real and simulated metagenomics samples for several choices of the sample composition, both in terms of species abundance and the similarity among the constitutive genomes.

### Experiments on complex eukaryotic genomes

#### Experimental results on real data with varying ploidy

All the assemblers were tested on real HiFi reads for rice (homozygous diploid), potato (heterozygous diploid), and wax apple (autotetraploid). Detailed statistics on these datasets and the calculation methods of evaluation criteria are provided in **Methods**.

First, the assembly contiguity was assessed by QUAST. **Figure 2A** shows the cumulative total size of contigs and the number of contigs that have a size in the range encoded by the color in the legend, for the rice dataset (top), the potato dataset (middle), and the wax apple dataset (bottom). Observe that on potato and wax apple data, HiCanu, hifiasm, HiFlye and Peregrine produced longer, more contiguous assemblies. In particular, hifiasm produced a larger proportion of contigs longer than 10Mbp (blue area in **Figure 2A**). Verkko produced a long assembly composed primarily of relatively short contigs. MECAT2 instead produced the shortest assemblies. In **Figure 2C**, we ordered the assembled contigs by size and computed the cumulative contig length for different thresholds of the NG value. Observe that on all three datasets, hifiasm achieved the best contiguity.

**Figure 2.**
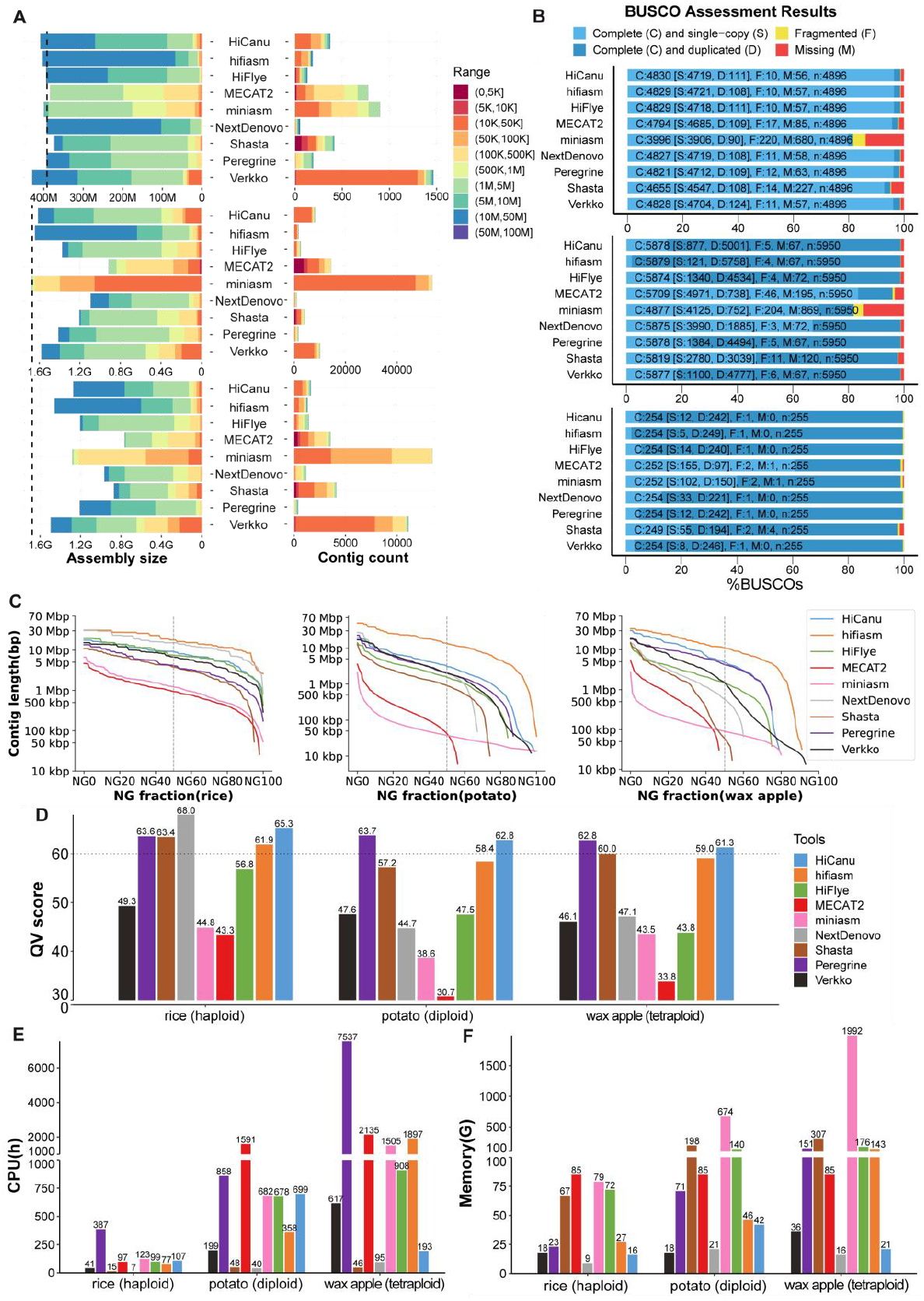
Summary of the performance of the genome assemblers on PacBio HiFi reads for rice (haploid), potato (diploid) and wax apple (tetraploid). (**A**) Cumulative total size of contigs (left) and number of contigs (right) that have a size in the range encoded by the color in the legend (top: rice; middle: potato; bottom: wax apple); the vertical dashed line indicates the expected genome size. (**B**) BUSCO completeness scores (top: rice; middle: potato; bottom: wax apple). (**C**) Contig length distribution for various choices of the NGx fraction threshold. (**D**) Quality value scores. (**E**) Running time analysis. (**F**) Memory usage analysis (legend on panel D also applies to panel E-F).

Second, the genome completeness was assessed by BUSCO. The set of conserved genes in *gramineae* (4896 genes in total) was used to assess the rice assemblies, the set of conserved genes in *cruciferae* (5878 genes) was used to assess the potato assemblies, and the set of *eukaryotic* conserved genes (5878 genes) was used to assess the wax apple. **Figure 2B** shows the BUSCO assessment results for rice (top), potato (middle) and wax apple (bottom). Observe that HiCanu, hifiasm, HiFlye, NextDenovo, Peregrine, and Verkko achieved over 98% completeness. According to this metric, miniasm did not perform well on rice and potato.

Third, the assembly accuracy was assessed using the QV score. The QV scores for all the assemblers on the three datasets are shown in **Figure 2D**. Observe that HiCanu, hifiasm, NextDenovo, Shasta, and Peregrine produced high QV across the three datasets. In contrast, MECAT2 and miniasm produced low accuracy assemblies on all datasets. Shasta generated poor accuracy assemblies on the wax apple, while the performance on rice and potato was satisfactory. It is noteworthy that Verkko had a worse QV score than HiCanu, despite the fact that it shares some of its codebase.

Fourth, CPU time and memory usage was collected. **Figure 2E** and **Figure 2F** show the run time and the memory usage for all assemblers on the three datasets. Observe that Shasta and NextDenovo consumed the lowest amount of resources. Peregrine was the slowest on rice and the wax apple datasets. MECAT2 was also slow, especially on the potato and the wax apple data set. NextDenovo and HiCanu used the smallest amount of memory. miniasm and MECAT2 used a much higher amount of memory, and the usage increased on higher-ploidy data sets.

Overall, hifiasm, HiCanu, HiFlye and Peregrine demonstrated a clear advantage over the other assemblers in terms of contiguity, completeness, and accuracy. In this group however, Peregrine required much higher computational resources, in particular related to CPU time. NextDenovo achieved good results on the rice datasets but did not perform equally well on the potato and wax apple genome.

#### Experimental results on synthetic datasets with varying ploidy

On synthetic dataset a more comprehensive evaluation of contiguity, completeness and accuracy was carried out due to the availability of the “ground truth” genome (i.e., one to four copies of the rice genome after introducing SNPs and structural variants; detailed statistics on these synthetic datasets are provided in **Methods**). **Figure 3** summarizes the experimental results on the nine HiFi assemblers on synthetic reads produced from the rice genome in haploid form (one copy of each chromosome), synthetic diploid (two copies) and synthetic tetraploid (four copies).

**Figure 3.**
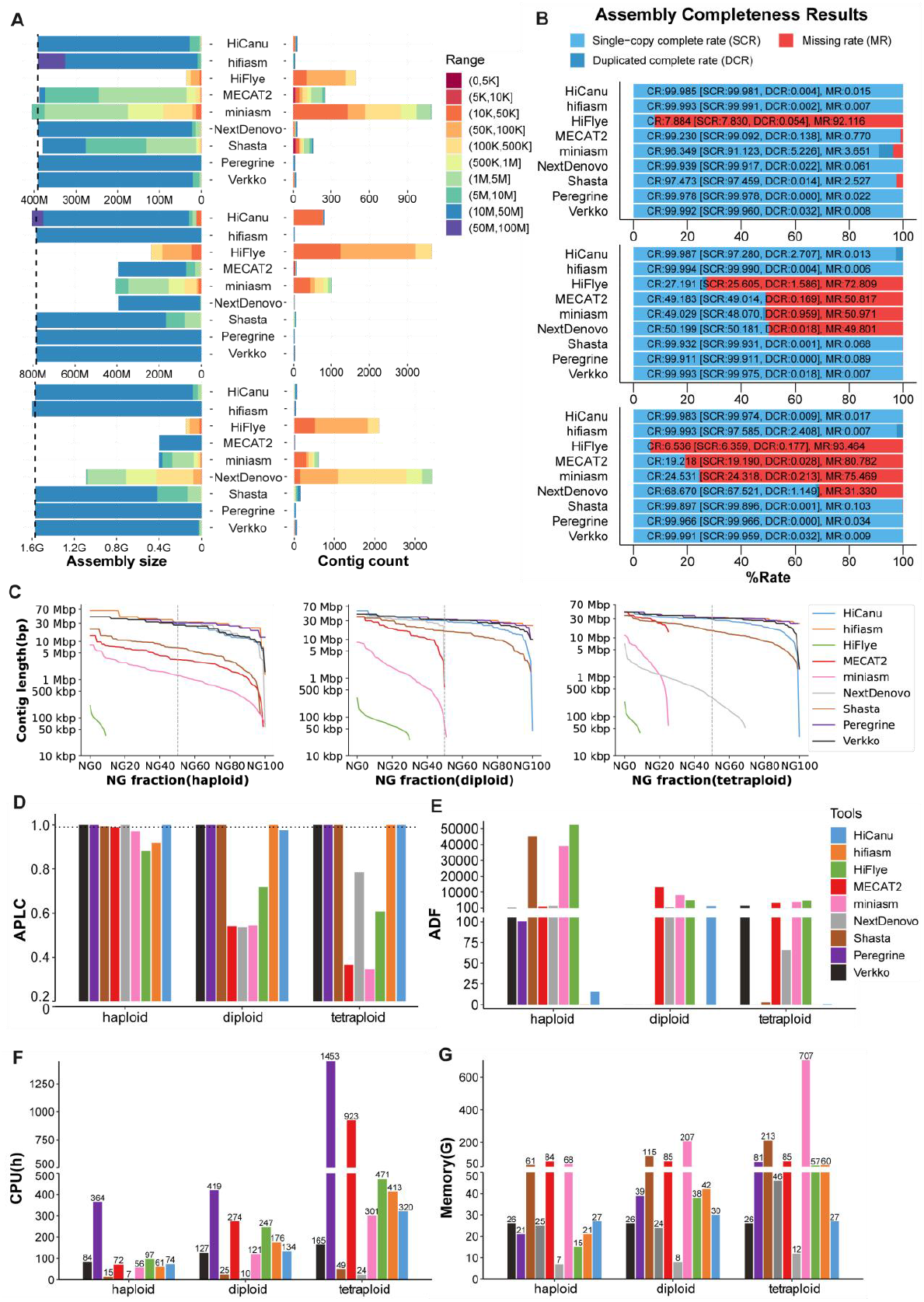
Summary of the performance of the genome assemblers on synthetic HiFi reads for rice in haploid form, synthetic diploid and synthetic tetraploid. (**A**) Cumulative total size of contigs (left) and the number of contigs (right) that have a size in the range encoded by the color in the legend (top: haploid rice; middle: synthetic diploid rice; bottom: synthetic tetraploid rice); the vertical dashed line indicates the expected genome size. (**B**) Single complete rate, duplicated complete rate and missing rate (top: haploid rice; middle: synthetic diploid rice; bottom: synthetic tetraploid rice). (**C**) Contig length distribution for various choices of the NGx fraction threshold. (**D**) Average proportion of largest category (APLC), the horizontal dashed line is APLC=0.99. (**E**) Average distance difference (ADF). (**F**) Running time analysis. (**G**) Memory usage analysis (legend on panel E also applies to panel F-G).

**Figure 3A** shows the cumulative total size of contigs and the number of contigs that have a size in the color-coded range, for the rice haploid dataset (top), the rice diploid dataset (middle), and the rice tetraploid dataset (bottom). Observe that hifiasm, Peregrine, HiCanu, Verkko and Shasta produced highly contiguous assemblies in all three datasets, with a high fraction of contigs longer than 10Mbp (blue sub-bars on the left of **Figure 3A**). HiFlye had the worst performance on all three synthetic datasets. A deeper analysis to explain HiFlye’s poor performance was conducted in the section **“HiFlye tested on synthetic datasets on varying sequencing error rates”**. miniasm and MECAT2 had an adequate performance only on the haploid data set, possibly because these assemblers were not designed to handle polyploid data sets. The results in **Figure 3A** were consistent with the NG curves in **Figure 3C**. In this latter figure, hifiasm, Peregrine, Verkko and HiCanu produced a higher NG50 on all synthetic datasets. **Supplementary Table 12** reports QUAST’s evaluation of the assemblies, including the number and length of mis-assembled contigs, the duplication ratio (which measures redundant contigs), the fraction of the genome covered by the assembly and the number of mismatches/indels in the assembly. Consistent with results above, hifiasm, Peregrine, Verkko and HiCanu achieved genome fraction higher than 99%, with duplication ratio very close to 1.0 and a low number of mismatches/indels and mis-assemblies. HiFlye, miniasm and MECAT2 did not perform as well.

In **Figure 3B**, we used the three-completeness metrics described in **Methods**, namely CR, SCR, and DCR, to evaluate the performance of the assemblers. HiCanu, Verkko and hifiasm had the best performance according to these three metrics, achieving a completeness rate of ∼99.9%. NextDenovo and MECAT2 performed well on haploid, but were inadequate on diploid and tetraploid. HiFlye had the worst performance again.

**Figure 3D** shows the average proportion of the largest category score (APLC), while **Figure 3E** summarizes the average distance difference scores (ADF). Both accuracy metrics were defined in **Methods**. HiCanu, Peregrine, Verkko and hifiasm performed well according to these criteria across varying ploidy, with APLC close to 1.0, and low values for ADF. On diploid and tetraploid datasets, other assemblers also had a small APLC. HiFlye and MECAT2 produced a high ADF which is consistent with the large number of misassembled contigs detected by QUAST for these tools (**Supplementary Table 12**).

As expected, time and memory usage on synthetic data was similar to the usage on real datasets (**Figure 3F**). Observe that Shasta and NextDenovo were the fastest on all synthetic datasets while HiFlye, MECAT2, and Peregrine were the slowest. Also observe in **Figure 3G** that NextDenovo used the smallest amount of memory, whereas miniasm, Shasta, and MECAT2 used the largest amount.

In summary, hifiasm, Peregrine, Verkko, HiCanu and Shasta produced assemblies with higher contiguity, completeness, and accuracy than the other assemblers on synthetic HiFi reads datasets across varying ploidy. NextDenovo had a good performance only in the haploid dataset. miniasm, NextDenovo and MECAT2 failed on the diploid and tetraploid datasets. HiFlye failed on all data sets; a deeper analysis will be carried out later.

#### Experimental results on synthetic datasets with varying sequencing coverage levels

An important quality of genome assembler is the ability to produce good assemblies even when the sequencing coverage is less than optimal. In this section we tested HiCanu, Verkko, hifiasm, and HiFlye using synthetic datasets from the synthetic heterozygous diploid genome with sequencing depths of 10x, 20x, 30x, and 50x. Detailed statistics on the synthetic datasets are provided in **Methods**. Experimental results are summarized in **Figure 4** and tabulated in **Supplementary Table 13**.

**Figure 4.**
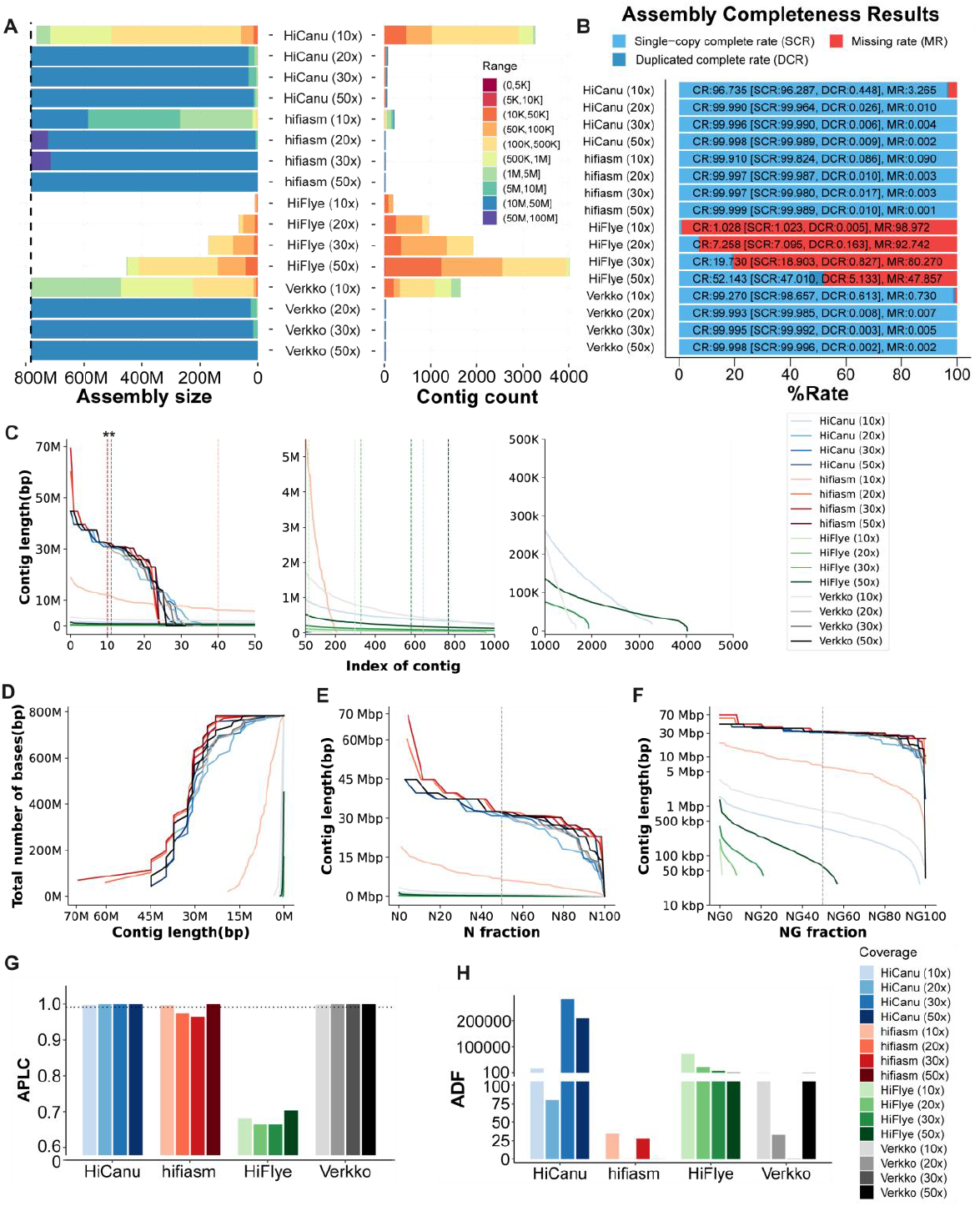
Summary of the performance of HiCanu, Verkko, hifiasm, and HiFlye on synthetic HiFi reads with coverage 10x, 20x, 30x and 50x. (**A**) Cumulative total size of contigs (left) and the number of contigs (right) that have a size in the range encoded by the color in the legend. (**B**) Single complete rate, duplicated complete rate and missing rate. (**C**) Sorted contig length (left: longest 50; middle: index 50-1000; right: index 1000-500). (**D**) Cumulative assembly size as a function of the minimum contig length allowed in the assembly. (**E**) Nx length (the dashed line denotes the N50). (**F**) NGx length (the dashed line denotes the NG50). (**G**) Average proportion of largest category (APLC); the horizontal dashed line is APLC=0.99. (**H**) Average distance difference (ADF).

Several observations are in order. First, irrespective of the assemblers, there was a significant improvement on the assembly contiguity when the coverage increased from 10x to 20x. HiCanu’s N50 increased from 362,820 bps to 30,836,250 bps; Verkko’s N50 increased from 816,272 bps to 32,114,812 bps; hifiasm increased from 6,168,309 bps to 32,623,532 bps. At the same time the ADF of all assemblers decreased when the coverage increased from 10x to 20x. Interestingly, Verkko’s ADF continued to decrease when coverage increased from 20x to 30x, but it was not the case for HiCanu and hifiasm. When the coverage increased from 10x to 20, NA50 and NGA50 improved about 80 times for HiCanu, 40 times for Verkko and 5 times for hifiasm. From 20x to 50x, the change of N50, NA50, and NGA50 were less pronounced.

**Supplementary Table 13** and **Figure 4B** show that when the coverage increased from 10x to 20x, the assembly completeness for HiCanu and Verkko increased dramatically, whereas when the coverage increased from 20x to 50x the assembly completeness did not change significantly. Also observe that the increase in coverage did not affect hifiasm’s assembly completeness, which was remarkably stable over different coverages. HiFlye had better completeness with increasing coverage, but overall HiFlye’s assemblies were much worse than the other three assemblers.

The accuracy assessment was based on APLC, ADF, mismatches per 100 kbp and number/length of mis-assembled contigs. The APLC for HiCanu increased with higher coverage. hifiasm was able to produce more contigs longer than 50 Mbp (purple color in **Figure 4A**) with 20x and 30x coverage, however **Supplementary Table 13** and the APLC indicates a mis-assembly on those long contigs. With 50x coverage, hifiasm corrected this mistake (**Supplementary Table 13** and **Figure 4G**). With increased coverage Verkko produced an improvement in ADF.

In summary, HiCanu and Verkko’s assemblies were more sensitive to the sequencing coverage, and they had an unsatisfactory performance at 10x. Instead, hifiasm had a more predictable and consistent performance across different choices of coverage, and its assemblies improved with increasing coverage.

#### Experimental results on synthetic datasets with varying heterozygosity rates

Another important quality of genome assembler is the ability to deal with various levels of heterozygosity in diploid (or polyploid) genomes. In this section, we used the synthetic heterozygous diploid genome and generated synthetic reads at 20x coverage, by varying the heterozygosity rates. Specifically, we used the SimSID script to introduce heterozygosity rates of 0.5%, 1.0%, 1.5% and 2.5% (see **Methods**). Detailed statistics for the synthetic datasets are provided in **Methods**. Experimental results for HiCanu, Verkko, hifiasm, and HiFlye on these data sets are summarized in **Figure 5** and tabulated in **Supplementary Table 14**.

**Figure 5.**
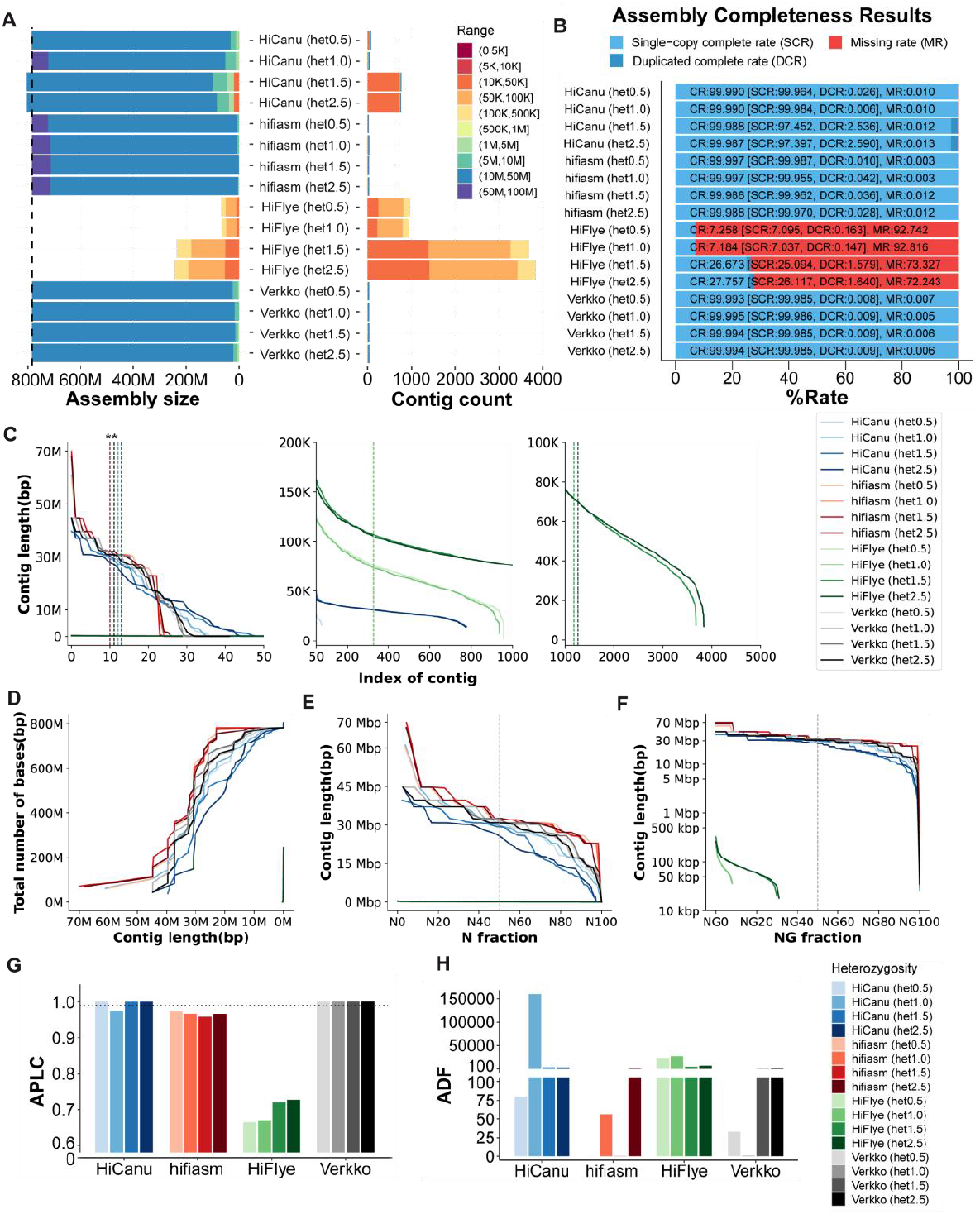
Summary of the performance of HiCanu, Verkko, hifiasm, and HiFlye on synthetic HiFi reads with heterozygosity rates 0.5%, 1.0%, 1.5% and 2.5%. (**A**) Cumulative total size of contigs (left) and the number of contigs (right) that have a size in the range encoded by the color in the legend. (**B**) Single complete rate, duplicated complete rate and missing rate. (**C**) Sorted contig length (left: longest 50 contigs; middle: index 50-1000; right: index 1000-500). (**D**) Cumulative assembly size as a function of the minimum contig length allowed in the assembly. (**E**) Nx length (the dashed line denotes the N50). (**F**) NGx length (the dashed line denotes the NG50). (**G**) Average proportion of largest category (APLC), the horizontal dashed line is APLC=0.99. (**H**) Average distance difference (ADF).

Several observations are in order. In terms of contiguity, **Figure 5A** shows that HiCanu’s assembly contiguity degraded with higher levels of heterozygosity while hifiasm’s had a stable performance across all datasets. In fact, hifiasm had a better performance on all metrics compared to the other assemblers (**Figure 5**). HiFlye again produced incomplete, small assemblies on all synthetic datasets. In terms of completeness (CR, SCR, DCR), hifiasm, HiCanu and Verkko had consistently good performance with varying heterozygosity rates.

With respect to assembly accuracy, there was no clear trend associated with increasing levels of heterozygosity. hifiasm’s number of mismatches per 100 kbp assessed by QUAST was less than 0.1 in all datasets (**Supplementary Table 14**). However, HiCanu’s APLC was better than hifiasm. hifiasm’s lower APLC was caused by a single mis-assembly on a 70Mbp contig (**Supplementary Figure 2**). hifiasm produced the highest accuracy assemblies based on the ADF metric.

In summary, while HiCanu and HiFlye’s performance degraded with increasing heterozygosity, Verkko and hifiasm has a consistently good performance across data sets. hifiasm, Verkko and HiCanu performed well on low heterozygosity datasets. hifiasm performed well also on high heterozygosity datasets.

#### HiFlye tested on synthetic datasets on varying sequencing error rates

As reported previously, HiFlye produced high-quality assemblies on the three real datasets, but unsatisfactory and inconsistent results on synthetic datasets. We hypothesized that HiFlye could be very sensitive to the sequencing error rate. To test this hypothesis, we started from (i) a real homozygous diploid rice genome, (ii) a synthetic heterozygous diploid genome and (iii) synthetic autotetraploid genome (see **Methods**), and generated synthetic datasets with different error rates using PBsim associate with the error models learned from on real HiFi datasets. The list of datasets used for learning the error models are shown in **Methods**. HiFlye’s results on these synthetic datasets are shown in **Supplementary Table** 1**5**.

As hypothesized, the assembly results of HiFlye decreased with higher sequencing error rates. Lower sequencing error rates led to improved completeness and contiguity in the assembly results. Specifically, at a read accuracy of 99.8474%, N50s and NA50s were consistently lower than 100 kb for different ploidy. However, at a significantly higher accuracy of 99.99%, HiFlye encountered memory issues when handling diploid and tetraploid genomes. However, the reason behind this memory limitation is unclear.

### Experimental results on metagenomic samples

For the metagenome assembly evaluation, we compared the performance of two metagenomic assemblers, namely hifiasm-meta (24) and metaFlye (27), against two general assemblers, namely HiCanu (19) and NextDenovo.

On real metagenomic samples, the quality of assemblies was evaluated in three steps. (i) The completeness and contamination of each assembled contig were assessed using CheckM (51), using marker genes that are specific to each species inferred lineage within a reference genome tree. (ii) CheckM and the Genome Taxonomy Database Tool Kit (GTDB-Tk) (52,53) were used to identify and classify the assembled bacterial contigs in a reference genome tree; assemblies were compared using the numbers of identified contigs at different taxonomic levels. (iii) The numbers of conserved 16s ribosomal RNA (rRNA) and 16s rRNA clusters were used to evaluate the completeness of the metagenomic assemblies; Bacterial and Archaeal Ribosomal RNA Predictor (Barrnap) was used to detect the 16s rRNA sequences, and VSEARCH (54) was used for clustering 16s rRNAs (using a minimum of 97% identity).

On synthetic data, the quality of assemblies was evaluated with MetaQUAST (40) by comparing them against the “ground truth” microbial genomes.

#### Experimental results of real metagenomic data

As mentioned above, the sheep gut metagenome data set (47) was used to test the four assemblers. A summary of the statistics of this data set is provided in **Methods**. Experimental results are illustrated in **Figure 6** and tabulated in **Supplementary Table 16**. Observe in **Supplementary Table 16** that hifiasm-meta produced the assembly with the largest total size, the largest number of contigs, the largest number of contigs longer than 1 Mbp, and the largest number of circular contigs longer than 1 Mbp. In addition, hifiasm-meta’s assembly contained twice or more 16s rRNAs compared to the other assemblies, and it identified the largest number of rRNA clusters. Observe in **Figure 6A** that hifiasm-meta produced the highest number of high-quality contigs, i.e., contigs with CheckM completeness rate higher than 90% and CheckM contamination rate lower than 10%. HiCanu produced the second highest number of contigs with a completeness rate larger than 90%. metaFlye generated a large number of contigs but with comparatively lower quality. In terms of contamination, metaFlye worked best with an average contamination rate lower than 1% (**Figure 6B**). Observe in **Figure 6C** that the cumulative length of hifiasm-meta assembly was about two times larger than the second largest one. metaFlye, HiCanu, and NextDenovo generated a similar numbers of long contigs (longer than 1 Mbp). While NextDenovo produced fewer short contigs, its N50 was significantly higher than the other assemblers (**Figure 6D**). To find out if the high N50 was solely due to the small number of short contigs, we sorted the contigs from the longest to shortest and drew the contig length distribution in **Figure 6E**. Observe that the curve for NextDenovo is always below those of the other assemblers, which confirms our hypothesis. hifiasm-meta’s curve is always above those of other assemblers, while metaFlye is in “second place”.

**Figure 6.**
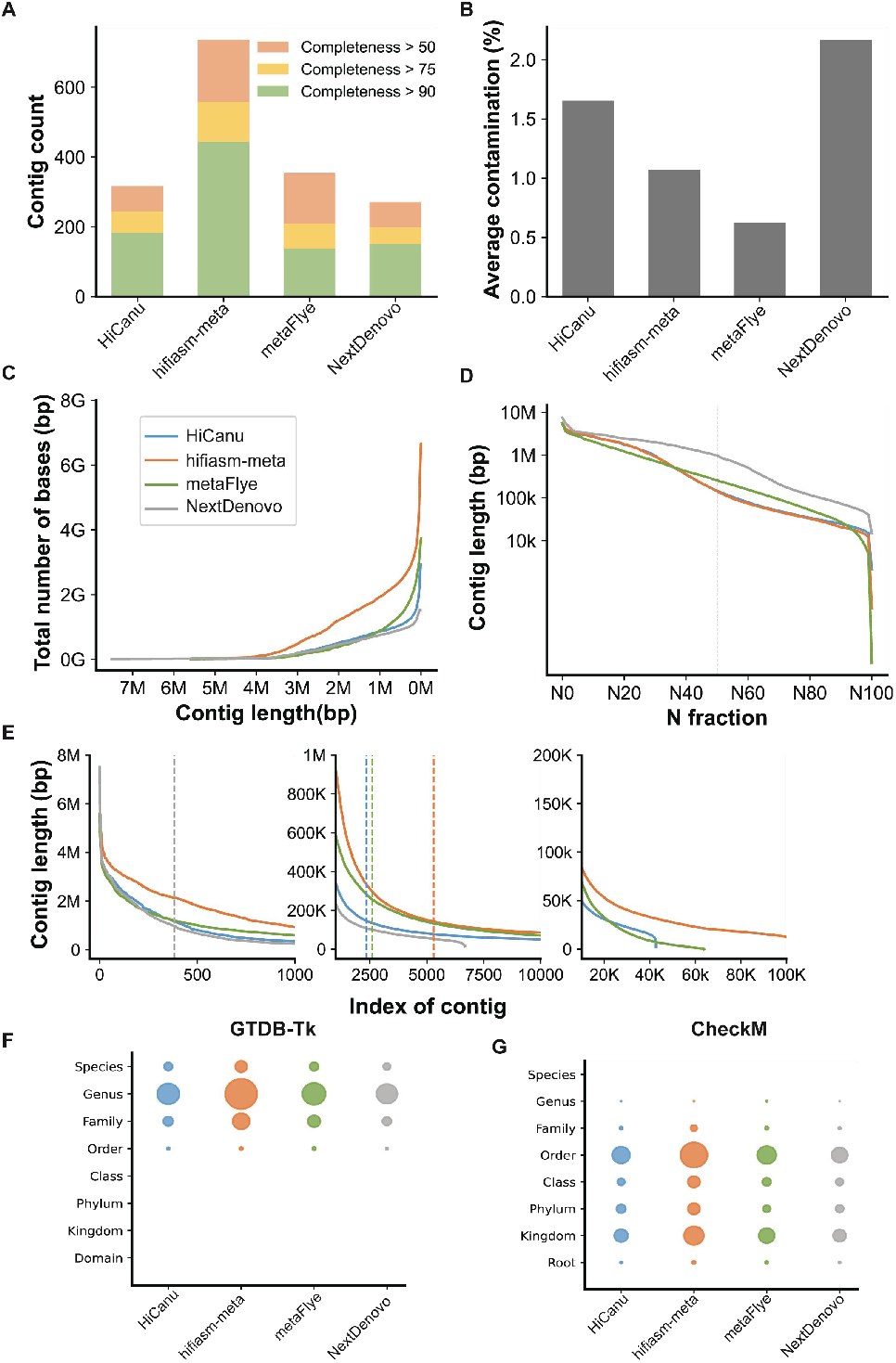
Summary of the performance of HiCanu, hifiasm-meta, metaFlye and NextDenovo on the sheep gut metagenome dataset. (**A**) Distributions of contigs in different levels of completeness; completeness was calculated by CheckM. (**B**) Average contamination rate calculated by CheckM. (**C**) Cumulative assembly size as a function of the minimum contig length allowed in the assembly. (**D**) Nx length (the dashed line denotes the N50). (**E**) Sorted contig length (left: longest 1000; middle: index 2500-10000; right: index 20k-100k). (**F**) Taxonomic classification of contigs longer than 1 Mbp with GTDB-Tk; the size of each circle represents the number of contigs identified at each taxonomic level. (**G**) Taxonomic classification of contigs longer than 1 Mbp with CheckM; the size of each circle represents the number of contigs identified at each taxonomic level.

To further compare the assemblies, we used CheckM in conjunction with GTDB-Tk to carry out the taxonomic identification and classification for the assembled contigs longer than 1 Mbp. Although the CheckM tends to classify contigs to higher taxonomic levels (mostly from Order to Kingdom) while GTDB-Tk tends to classify to lower ones (mostly Species to Family), they both indicate that hifiasm-meta is capable of assembling the largest number of species. Details of this experiment are shown in **Supplementary Figure 3**.

#### Experimental results of synthetic metagenomic data

Next, we tested the four assemblers on the synthetic dataset described in **Methods**. Completeness, contiguity and accuracy of the assemblers were evaluated using MetaQUAST and the five additional criteria described in **Methods. Figure 7** illustrates the experimental results. Observe in **Figure 7B** that all assemblers achieved low single-copy completeness rate (SCR) and duplicated completeness rate (DCR) (lower than 50%), which may be due to the non-uniform abundances of species in the sample. hifiasm-meta produced the most contiguous and complete assembly among all the tools, followed by metaFlye and HiCanu. HiCanu and hifiasm generated assemblies with better accuracy than other two assemblers, including higher average proportions of the largest category (APLC), lower average distance differences (ADF), lower misassembled rates, lower numbers of mismatches per 100 kbp, and lower numbers of indels per 100 kbp (**Figure 7** and **Supplementary Table 17**).

**Figure 7.**
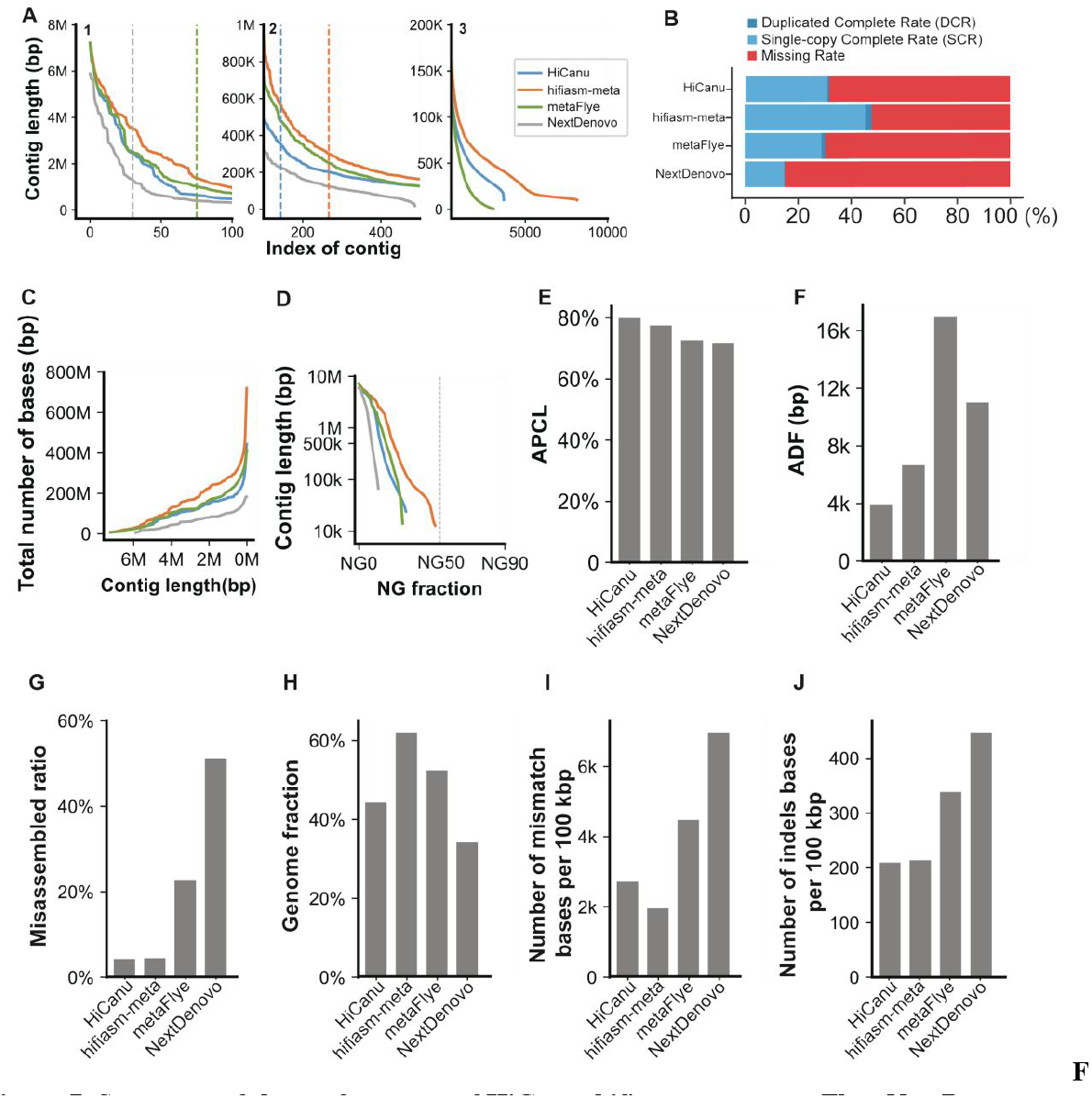
Summary of the performance of HiCanu, hifiasm-meta, metaFlye, NextDenovo on a synthetic dataset. (**A**) Sorted contig length (left: index 1-100; middle: index 1-500; right: index 1-10,000), the dashed line represents the N50 contig length. (**B**) Single-copy completeness rate (SCR) and duplicated completeness rate (DCR). (**C**) Cumulative assembly size as a function of the minimum contig length allowed in the assembly. (**D**) NGx length (the dashed line denotes the NG50). (**E**) Average proportion of largest category (APLC). (**F**) Average distance difference (ADF) (**G**) Misassembled contig rate computed by MetaQUAST. (**H**) Genome fraction computed by MetaQUAST. (**I**) Number of mismatches per 100 kbp computed by MetaQUAST. (**J**) Number of indels per 100 kbp computed by MetaQUAST.

#### Comparisons on different taxonomic levels

To compare the performance of the two metagenomic assemblers (hifiasm-meta and metaFlye) we tested them on several synthetic datasets (described in **Methods**) for various choices of the sequence similarities.

Experimental results in **Figure 8B and Supplementary Table 18** show that assemblies produced by hifiasm-meta had higher completeness (i.e., the genome fraction computed by MetaQUAST) than metaFlye’s assemblies. Also observe that the completeness of metaFlye’s assemblies declines faster than hifiasm-meta’s, as the similarity of genomes increases. However, hifiasm-meta assemblies had a genome fraction larger than 100% which indicated more redundant sequences than metaFlye. This redundancy was also shown in the unique k-mer based completeness criteria (**Figure 8G**).

**Figure 8.**
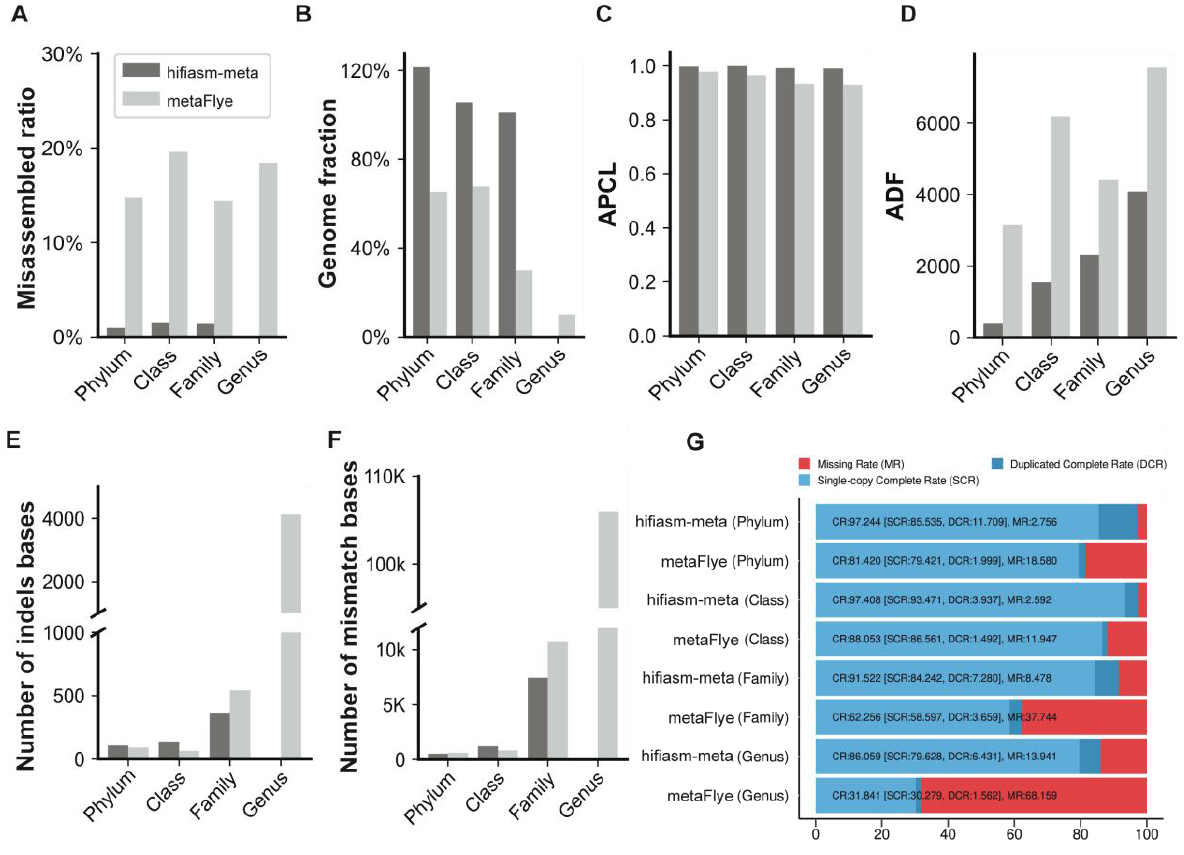
Comparison of hifiasm (dark gray) and metaFlye (light gray) on dataset at the phylum, class, family and genus levels. (**A**) Misassembled contig rate from MetaQUAST. (**B**) Genome fraction from MetaQUAST. (**C**) Average proportion of largest category (APLC). (**D**) Average distance difference (ADF). (**E**) Number of indels per 100 kbp from MetaQUAST. (**F**) Number of mismatches per 100 kbp from MetaQUAST. (**G**) unique *k*-mer based complete rates (including single complete rate, duplicated complete rate and missing rate). At the Genus level, MetaQUAST was not able to generate evaluation results for the hifiasm-meta, leading to the missing bars in the figures.

**Figure 8A** and **Figure 8C-F** show that hifiasm-meta’s assemblies are more accurate than metaFlye. As the sequence similarity of genomes increases, the accuracies of hifiasm-meta and metaFlye both decline sharply in terms of ADF, the number of indels per 100 kbp and the number of mismatches per 100 kbp. However, for both hifiasm-meta and metaFlye, APLC does not change significantly at different taxonomic levels.

## DISCUSSION

As mentioned above, choosing the “best” assembler to assemble a new genome is a daunting proposition, because the performance of an assembler often depends on the genome ploidy, repetitive content, size, and heterozygosity, and many other factors. To the best of our knowledge, there are very few comprehensive studies that can guide users on the choice of the most appropriate assembler for their data. Even if we focus only on HiFi data, we are not aware of any comprehensive study that compares the performance of all modern HiFi assemblers on large eukaryotic genomes. As mentioned in the Introduction, studies (37) and (38) are exclusively focused on the yeast, drosophila and human genomes.

In this paper we addressed this shortcoming by carrying out an extensive set of experiments to assess the performance of the most popular genome assemblers for HiFi reads. Several real and synthetic datasets for complex eukaryotic genomes and metagenomes were used to test the contiguity, completeness, and accuracy of eleven assemblers. We hope that our data sets (which include HiFi reads for a newly sequenced tetraploid, the wax apple) will become a benchmark for future assembler development. Five novel *k*-mer-based criteria were introduced to help assess genome assemblies’ quality.

Our concluding remarks are in order. Overall, hifiasm performed consistently well across all experiments with varying ploidy, coverage and heterozygosity. hifiasm was also the least sensitive to the sequencing coverage, as long as it was higher than 20x. It clearly ranked first in the overall performance.

HiCanu also produced good assemblies, albeit not as good as hifiasm. Overall, it ranked second among all the assemblers we tested. On real data, HiCanu produced assemblies of a quality similar to hifiasm. While HiCanu’s accuracy was as high as hifiasm, HiCanu’s contiguity was often lower than hifiasm. Similar results were obtained on synthetic datasets.

Surprisingly, the performance of Verkko was not as good as HiCanu despite the fact that they share a similar codebase. Verkko produced assemblies with lower N50, lower largest contig and lower QV. We speculate that the difference is due to the different data structure used by Verkko (de Bruijn graph) compared to HiCanu (overlap graph).

Unfortunately, HiFlye failed on all simulated datasets. Our experiments seem to indicate that HiFlye is very sensitive to the sequencing error rate of the input reads. Surprisingly this mediocre performance seems to contradict the results in (37), which show that HiFlye outperformed HiCanu, hifiasm, NextDenovo and Flye on HiFi reads for baker’s yeast. Three factors could explain the difference between our experimental results and (37). First, HiFlye might work better on the yeast genome because of its relatively small size and small amounts of repetitive content. In our experiments, HiFlye worked better on the rice genome than more complex genomes such as potato and wax apple. Second, the yeast HiFi data used in (37) might have had a lower sequencing error rate compared to ours. Our experiments show that HiFlye is extremely sensitive to sequencing errors. Third, (37) used a reference yeast genome that is not the same strain as the one sequenced with PacBio HiFi, which could have led to evaluation inaccuracies.

Shasta and Peregrine had good performance across all ploidy on real datasets. Their main limitation is the high resource consumption both in time and memory occupation. NextDenovo and MECAT2 only showed good performance in haploid datasets. On real datasets, miniasm produced genome assemblies of a size similar to hifiasm, but had lower contiguity and lower accuracy, mainly with respect to BUSCO and QV.

hifiasm-meta performed consistently well on real metagenomic data and synthetic meta-genome data sets at different taxonomic levels. NextDenovo had the poorest performance in terms of genome size, contiguity, and accuracy (e.g. highest contamination rate) on the real metagenomic data sets. While HiCanu is not specifically designed for metagenome assembly, it performed comparably well to metaFlye, particularly in synthetic datasets, achieving higher accuracy and completeness. metaFlye outperformed HiCanu and NextDenovo in real metagenomic data, still exhibited a significant disparity in terms of completeness and contiguity compared to hifiasm-meta. Overall, none of the assembled genomes achieved a satisfactory NG50 on synthetic datasets.

In the last twenty years, breakthrough algorithmic advances in genome assembly have enabled scientists to assemble larger and more complex eukaryotic genomes. Despite these advances, our experiments show that there is still space for improving genome assemblers. For instance, our experimental results show that none of the assemblers was able to generate chromosome-level (or telomere-to-telomere) assemblies solely from HiFi reads. More efforts are needed to improve the assemblies’ contiguity, as well as the assembly quality in low-coverage and repetitive regions (e.g., centromere and ribosomal DNA).

The problem of assembling genomes from metagenomic samples is even more difficult. Our experiments clearly highlighted the challenges of assembling many genomes simultaneously. For instance, hifiasm-meta often generated redundant assemblies, while metaFlye produced assemblies with low completeness. No metagenome assembler performed well on microbial metagenomic data sets with low coverage, leading to low completeness and uneven abundances. New methods are needed for improving the assembly quality of low-coverage genomes.

Finally, we believe that assembly evaluation methods are currently insufficient. BUSCO is a great tool, but it falls short when evaluating the completeness of eukaryotic assemblies when those species contain few conserved genes. QUAST is very useful, but cannot be used to evaluate the assembly quality for repetitive regions due to unreliable sequence alignments. For metagenomic assembly evaluations, CheckM can provide misleading outcomes. For example, it can give a good score to an assembly that contains a mix of sequences from closely-related genomes. We believe that new evaluation methods are needed for more accurately evaluating genome assemblies.

## CONCLUSION

In this study, we carried out a comprehensive benchmarking of eleven assemblers on eukaryotic genomes and metagenomes. On eukaryotic genomes we measured contiguity, completeness, and accuracy across varying ploidy, coverage levels, and heterozygosities. On metagenomes, we measured contiguity, completeness, and accuracy across different composition profiles and different taxonomic levels. Our experiments clearly show that hifiasm and hifiasm-meta should be the first choice for assembling eukaryotic genomes and metagenomes with HiFi data.

## DATA AVAILABILITY

### Data availability

The synthetic datasets generated as part of this study are available in the Agricultural Genomics Institute at Shenzhen repository, ftp://ftp.agis.org.cn/~panweihua/benchmark/. The real wax apple dataset generated is available in NCBI website (see **Supplementary Table 3** for details).

### Code availability

The script that implements the five quality criteria can be obtained at https://github.com/rookieluohh/benchmark.

## AUTHOR CONTRIBUTIONS

W. Yu designed the experiments and wrote the initial draft of the manuscript. H. Luo proposed the five criteria and carried out some genome assemblies. J. Yang designed the experiments for the metagenomic study and performed the metagenomic assembly. S. Zhang prepared the simulated datasets. H. Jiang supported H. Luo for eukaryote genome assembly. X. Zhao performed genome assembly and the outcome assessment. X. Hui and D. Sun performed metagenomic assembly evaluation. L. Li prepared the samples for the wax apple and conducted data quality control. X. Wei provided the wax apple materials. S. Lonardi edited the manuscript and supervised the project. W. Pan supervised the project and edited the manuscript.

## ACKNOWLEDGEMENTS

We thank Yongyao Li from Agricultural Genomics Institute at Shenzhen, [Chinese Academy of](https://www.caas.cn/en/) Agricultural Sciences, for his technical support in data storage and maintenance. We also thank the funding support for this project.

## CONFLICT OF INTERST

The authors declare that they have no competing interests.

